# Buffering role of HSP shapes the molecular evolution of mammalian and human genomes at short and long-term scales

**DOI:** 10.1101/2022.11.11.516130

**Authors:** Valeriia Timonina, Evgenii Tretiakov, Andrey Goncharov, Konstantin Gunbin, Jacques Fellay, Konstantin Popadin

## Abstract

Heat shock proteins in parallel with their main and originally discovered function – maintenance of folded proteins under stressful conditions, can play also background buffering role – by folding proteins with an excess of slightly-deleterious nonsynonymous variants (SDNV). Here we tested several scenarios of this buffering role. On the comparative species scale, we demonstrated that low-Ne species are characterized by a higher expression level of hsp90 which can be explained by the excess of SDNV. On the comparative tissue level, we showed that long-lived tissues have also a higher hsp90 expression level, which can be advantageous to maintain the functionality of proteins. On the comparative gene level, we demonstrated that purifying selection of hsp90 in low-Ne-species did not relax as strongly as it happens for control genes, similar to hsp90. Additionally, we demonstrated that hsp clients versus non-clients are characterised by decreased level of selective constraints; demonstrate stronger relaxation of purifying selection in low-Ne species; have an excess of slightly-deleterious variants associated with complex disease phenotypes in humans; have an excess of pathological variants associated with clinical phenotypes in humans, suggesting that clients, being buffered by hsp90 can degenerate a bit more as compared to non-clients. Altogether, our results show that the secondary role of hsp, buffering of SDNV, is widespread and universal affecting properties of species, tissues and genes. A deep understanding of the buffering role of hsp90 will help to predict the deleterious effect of each variant in the human genome more precisely as well as will extend the application of the effectively-neutral theory of molecular evolution.

## INTRODUCTION

Fixation of slightly deleterious variants due to genetic drift is accelerated in species with low effective population size (Ne) (Popadin et al. 2007; Ohta 1973). Thus, it is expected that genomes of such species carry an increased burden of suboptimal variants, including the nonsynonymous ones, which may decrease the fitness (Popadin et al. 2007) and even lead to extinction due to mutational meltdown. Here, we hypothesize that heat shock proteins and especially, hsp90, can play a buffering role of a compensatory mechanism decreasing the deleterious effect of the mutational burden. Numerous experimental and theoretical works unambiguously show a compensatory effect of hsp90 on short evolutionary intervals in humans (“HSP90 Shapes the Consequences of Human Genetic Variation” 2017a), drosophila (Rutherford and Lindquist 1998), etc, and in this work, we extend the time interval, potentially affected by the hsp90. In order to do it, we analyzed several evolutionary properties of hsp and its clients (protein-coding genes, frequently folded with the assistance of hsp) on both comparative species and human population genetic levels.

## RESULTS

### 1. The expression level of HSP90 is higher in primate species with low effective population size (Ne)

The expression level of hsp90, if essential for compensation of the increased mutational burden of low-Ne species, is expected to be higher in low-Ne species. To test it, we analysed an expression dataset (Rutherford and Lindquist 1998; Brawand et al. 2011) containing uniformly processed data for six tissues in 6 primate species with different Ne (Methods). We focused on hsp90 (HSP90AB1, ENSG00000096384) which is constitutively expressed in all mammals and which demonstrated a buffering role against deleterious variants in humans (“HSP90 Shapes the Consequences of Human Genetic Variation” 2017b), drosophila (Rutherford and Lindquist 1998) and other species.

Among analysed six primate species, great apes are characterised by significantly lower Ne (human: 13’100-16’200; gorilla: 28’400-56’900; chimp: 30’900-61’800; orangutan: 42’300-84’600, (Prado-Martinez et al. 2013)) as compared to Macaca mulatta (80’000, (Yuan et al. 2012)). Comparing the tissue-specific hsp90 expression level between the great apes and Macaca we indeed observed in 5 out of 6 tissues (all except testis) an increased hsp90 expression level in formers (paired by tissue Mann-Whitney U-test, Hochberg adjusted p-values when comparing with M. mulatta for H. sapiens - 0.01, P. paniscus - 0.02, P.troglodites - 0.1, G. gorilla - 0.005, P. pygmaeus - 0.01). Thus our first result is in line with the hypothesis that the hsp90 role (i.e. expression level) can be more important in species with low Ne.

### 2. The expression level of HSP90 is higher in tissues with increased cellular longevity (a low turnover rate)

Mammalian species with low Ne usually have a longer generation length. Thus the increased expression level of hsp90 in great apes can be advantageous due to both the necessity to compensate for the high mutational burden due to their low Ne and the necessity to maintain proteins in a functional state for an extended period due to their higher longevity. To test the effect of longevity separately, we analysed expression levels of hsp90 in human tissues with different turnover rates (hereafter, “cellular longevity”).

We performed a linear regression analysis using expression levels of hsp90 from 39 human tissues and the cell turnover rates collected for the corresponding tissues (Methods). We observed a significant positive correlation between the hsp90 expression level and the log10 of the tissue-specific turnover rates in days (slope = 9.427, p-value = 0.0374, R_sq_adj = 0.08784, N=39, Fig. 1A). Thus, long-lived tissues with a turnover rate taking many days, tend to have an increased expression level of hsp90 as compared to short-lived ones. To prove the robustness of this result, we additionally clustered all tissue into three groups by their turnover rate - fast-(uterus, colon/rectum, stomach, cervix, esophagus, head/neck, lymphoid and myeloid tissues), intermediate-(breast, prostate, skin, bladder, biliary, pancreas, liver, kidney) and slow-replicating (thyroid, lung, bone/soft tissue, CNS, ovary). We observed a significant difference between fast- and slow-replicating tissue (Fig. 1B, Mann-Whitney U-test p-value = 0.003), as well as between slow- and intermediate-replicating tissues (Mann-Whitney U-test p-value = 0.008), while we didn’t observe a significant difference between intermediate- and fast-replicating tissues (Mann-Whitney U-test p-value = 0.71).

**Figure 1.**
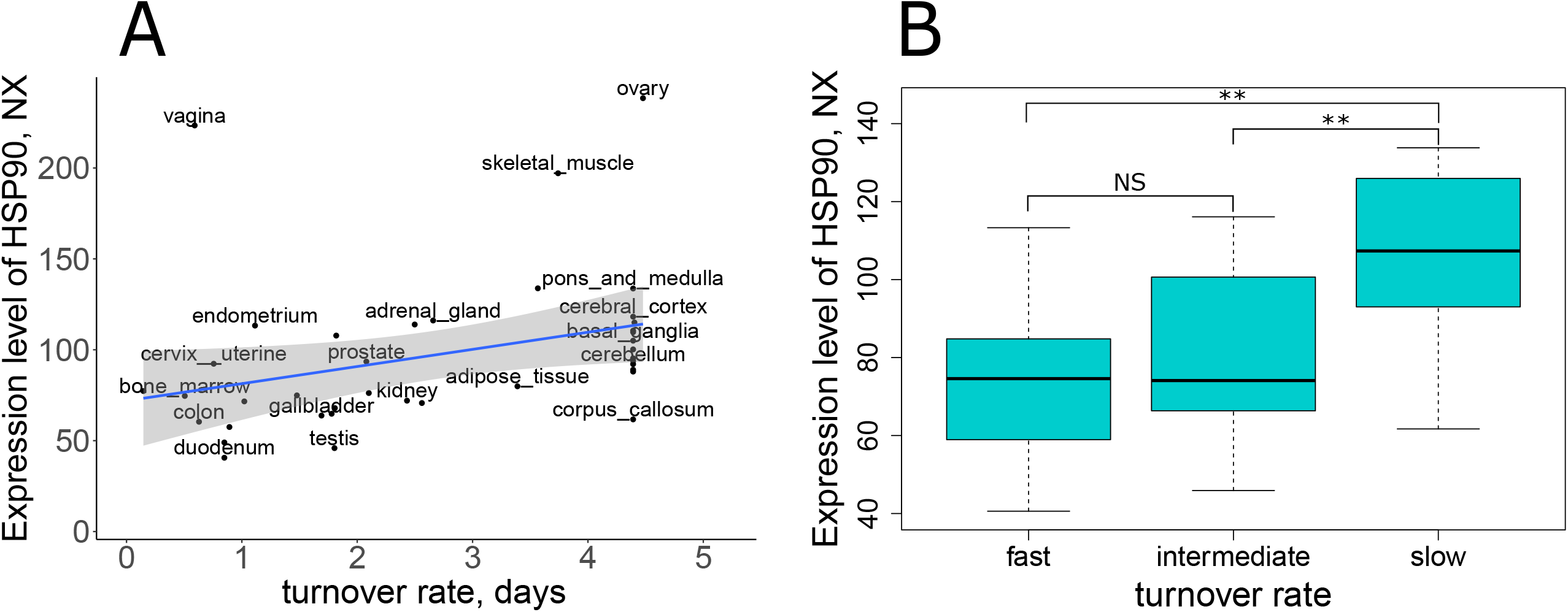
Hsp90 expression level is higher in tissues with a low turnover rate. An increased expression level of HSP90 in different tissues with slow turnover rates is demonstrated by (A) an ordinary linear model (slope = 9.427, p_value = 0.0374, R_sq_adj = 0.08784) and (B) a Box plot depicting hsp90 expression values in tissues, grouped by turnover rates: fast-, intermediate and slow-replicating tissues. Mann-Whitney U-test. Outliers are removed from (B). Significance code: ** - p-value < 0.01, NS - not significant.

These results show that the hsp90 role, approximated by its expression level, can be a function of both effective population size and longevity. Future studies are needed to deconvolute these factors.

### 3. Hsp90 demonstrates a strong purifying selection in species with low Ne

The functional importance of *hsp90* in species with different Ne can be tested not only by its expression level but also by its pattern of molecular evolution. It is known that species with a low effective population size accumulate more nonsynonymous substitutions as compared to species with a high effective population size. The commonly used approach to validate this statement is to correlate the species-specific rate of accumulation of nonsynonymous substitutions relative to synonymous substitutions, dN/dS, which is a proxy for selective constraints (the higher dN/dS the less stringent the purifying selection) with life-history traits, such as generation length, which approximates Ne (the higher the generation length the lower the Ne). Linear regression models of gene-specific dN/dS versus generation length typically show a positive slope, the magnitude of which can be interpreted as a gene-specific relaxation of purifying selection in species with low Ne. Assuming the unique buffering role of hsp90 in species with low Ne, we expect that hsp90, compared to other genes, shouldn’t relax intensely in such species. To test this, we performed a simple linear model of the dN/dS as a function of log10 generation length for 71 mammalian species. We derived the species-specific dN/dS values using codeml program from the PAML v4.9 package based on codon alignments from OrthoMaM v10c and phylogenetic tree from TimeTree (see Methods). Additionally, we retrieved the generation length of each mammalian species from the mammalian database (Marco et al. 2013).

Next, to compare the hsp-specific slope (relaxation) with other control genes we generated a subset of genes, similar to hsp90 and estimated for each such gene the level of relaxation of purifying selection using the above-described approach (see Methods). To generate such a control subset of genes we took into account 7 characteristics of evolutionary constraints (selective effects of heterozygous protein-truncating variants, probability of being intolerant to loss of function mutation, functional indispensability score, residual variation intolerance score, genome-wide haploinsufficiency score, Kn/Ks compared to mouse, branch) as well as 5 structural characteristics (gene length, number of transcripts, average exon length, average number of exons per transcript, GC-content) of ∼11500 human protein-coding genes. Taking all these parameters into account, we performed PCA and chose 300 genes closest to HSP90 in the space of the first and the second principal components (Fig 2A).

**Figure 2.**
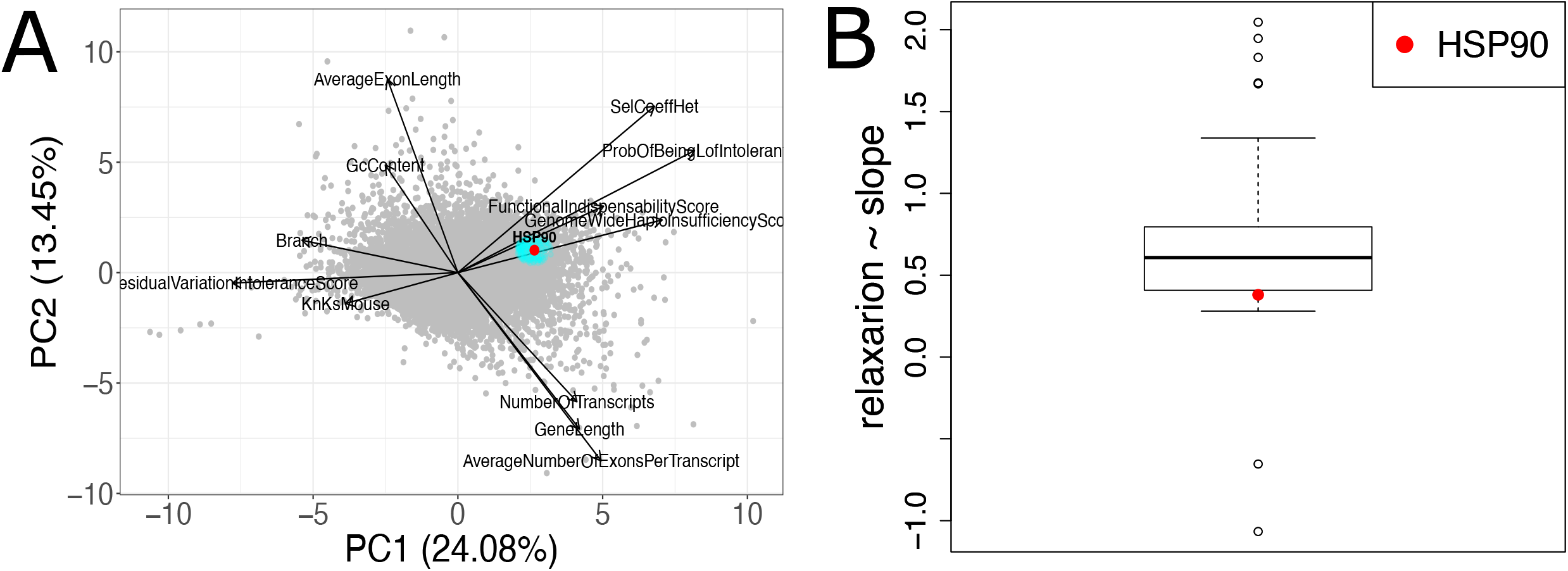
The strength of the purifying selection of HSP90 doesn’t relax strongly in species with low effective population size. A - Principal component analysis (PCA) plot depicting distance between ∼11500 human protein-coding genes and HSP90. Seven characteristics of evolutionary constraints and 5 structural characteristics (See description in text) were indicated as arrows (I found only 11 arrows?!! + not all text under arrows is readable). B - Box plot, showing nominally significant slopes (the level of relaxation) distribution from linear model Ensembl-derived Kn/Ks ∼ generation length (one-sample Mann-Whitney U-test, p-value < 0.001).

For selected control genes as well as for hsp90, we estimated the level of relaxation as a slope in the linear regression analysis of species-specific Ensembl-derived dN/dS (using 1-to-1 orthologs from Ensembl Compara) and log10 of species-specific generation length. In this approach, hsp90 relaxation was slightly higher than zero, however, when compared with controls (all nominally significant slopes) we still observed that hsp90 is in the lower quartile of the distribution of significant slopes (Fig 2B). This observation further supports that relaxation of purifying selection is stronger for control genes as compared to *hsp90, which is constrained due to its additional role of buffering the mutational burden*.

### 4. The human HSP90 clients are enriched in the low-constrained genes

It is expected that hsp clients, due to the compensatory role of hsp, can have an increased rate of accumulation of suboptimal amino acid substitutions ((Agozzino and Dill 2018; Alvarez-Ponce, Aguilar-Rodríguez, and Fares 2019), but see (Victor et al. 2020)). To test it we compared the evolutionary constraints of clients versus non-clients among the human protein-coding genes using principal component analysis (PCA). The PCA was based on gene structure features like Average exon length, GC content, Gene length, Number of transcripts, and an Average number of exons per transcript, as well as evolutionary constraints scores such as Residual Variation Intolerance Score, Branch, Genome-Wide Haploinsufficiency Score, Probability of being loss-of-function intolerant, Functional Indispensability Score, Selective effects for heterozygous protein-truncating variants (*SelCoefHet*), oe-lof-upper-bin, p (see Methods). We performed Principal Component Analysis (PCA) to take all these metrics into account simultaneously.

We observed that traits with high loadings on PC1 are relevant to evolutionary constraints: Residual Variation Intolerance Score, Branch, Genome-Wide Haploinsufficiency Score, Probability of being loss-of-function intolerant, Functional Indispensability Score, Selective effects for heterozygous protein-truncating variants (*SelCoefHet*), oe-lof-upper-bin, p. Thus we interpret the PC1 as the level of selective constraints (which is increased along the PC1). Traits with high loadings on PC2 are structural: Average exon length, GC content, Gene length, Number of transcripts, and an Average number of exons per transcript.

We found that clients of hsp90 versus all other genes are shifted left along the PC1 (Fig.3A bottom panels, p-value=0.01 Mann-Whitney U-test for PC1 comparing HSP90 (N=229) clients to the background (grey points, N = 11187)), towards the less constrained genes. We didn’t observe any significant shifts in PC2 (p-value=0.26). Thereby hsp90 clients demonstrated indeed a decreased level of selective constraints which is in line with our expectation of an increased rate of accumulation of suboptimal amino acid substitutions.

**Figure 3.**
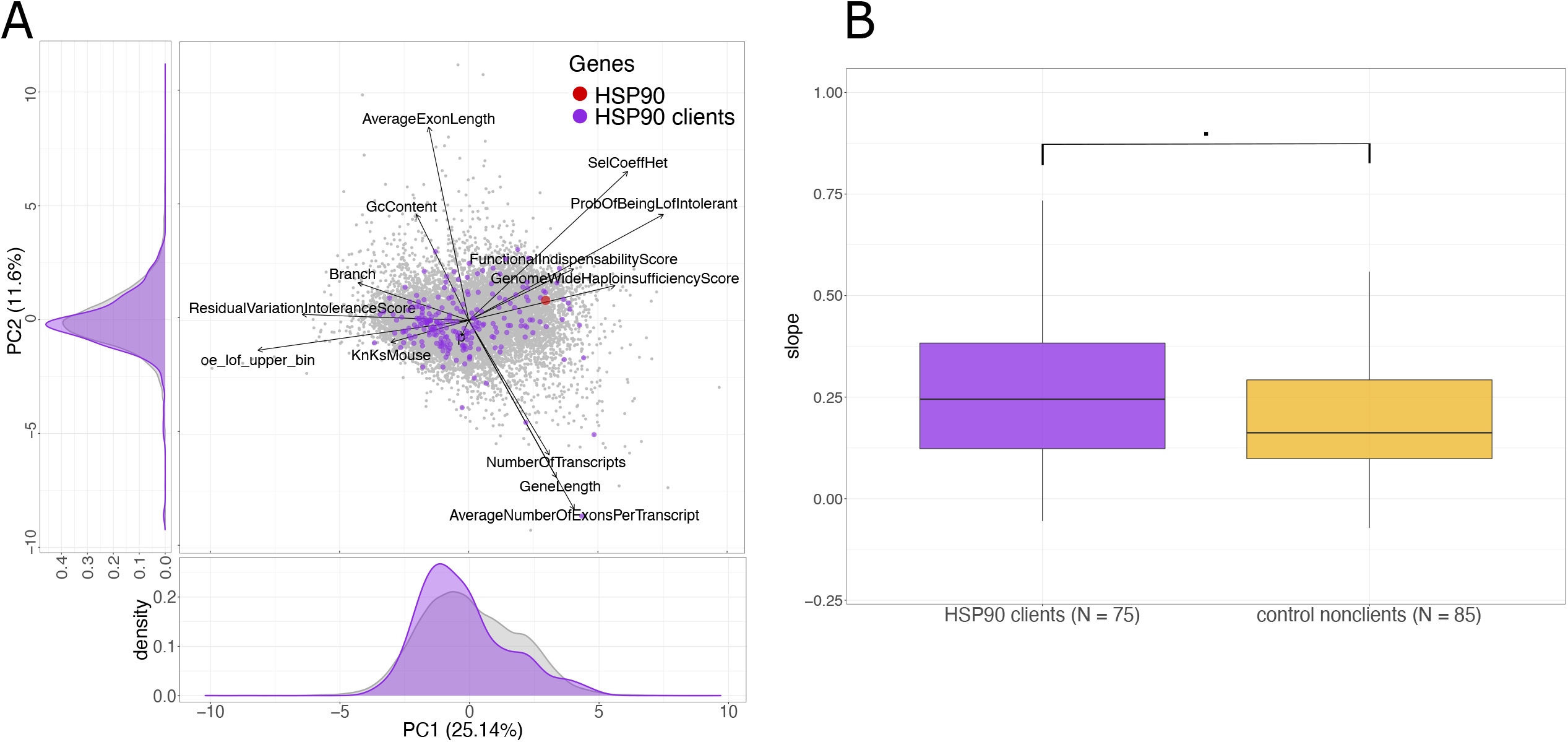
A - Projection of HSP90 clients to the first and the second Principal Components (PC1 and PC2) of Principal Component Analysis built by taking into account metrics of evolutionary constraint and structure of ∼11500 human protein-coding genes. Clients of HSPs are shifted on PC1 to the side of less constrained genes. B - Values of slopes for HSP90 clients and nonclients, genes, obtained by linear model dN/dS vs. log10 generation length (one-sided Mann-Whitney U-test, p-value = 0.06).

### 5. Purifying selection of hsp90 clients is more relaxed than that of nonclients

Clients are proteins whose foldings are supported by HSP activity. Because HSP90 can buffer the manifestation of deleterious mutations we expect its clients to accumulate more such mutations compared to non-clients. The first line of evidence we observed in the PCA (Fig.3A). To validate this result additionally we compared the level of relaxation of HSP90 clients (the client’s list was taken from Sahni N. *et al*., 2015) and control nonclient genes (the list of nonclients was derived from PCA as genes similar to clients in evolutionary constraints and structural characteristics, see Methods). We built the same linear model (species-specific paml-derived dN/dS vs. log10 generation length) as in the previous analysis with a correction for the phylogenetic relationship of species using PGLS (Phylogenetic generalized least squares) and compared the slopes of the model for clients and non-clients. We observed a marginally significant trend that follows our expectations - HSP90 clients have an increased slope (relaxation) as compared to the control nonclient genes (Fig.3B, *one-sided Mann-Whitney U-test, p-value = 0*.*06*).

As the buffering role of HSP90 was demonstrated previously from a comparative species analysis, we focused next on the potential consequences of the HSP90 buffering role for the human genome. We analysed the level of selective constraints (Chapter 5), integral mutational burden (Chapter 6) of clients versus nonclients and the probability of carrying pathogenic variants by clients and non-clients (Chapter 7).

### 6. Hsp90 clients have an increased burden of slightly-deleterious variants associated with complex human diseases

We considered variants associated with complex human diseases to approximate a mutational burden of slightly-deleterious variants segregating in different genes. We chose polygenic scores data for disease-associated phenotypes from PGS Catalog (https://www.pgscatalog.org) and extracted variants from 100kb intervals from the gene start for hsp90 clients and their control nonclients. Using this dataset, we compared the total number of variants associated with each phenotype, the number and proportion of positively associated variants, and the effect size of the associated variants. We observed a weak but highly significant difference in the proportion of variants with a positive effect on disease phenotype (Mann-Whitney U-test, p-value=1.8e-07, the median is 1.02 times higher in clients,) and the mean effect of variants (Mann-Whitney U-test, p-value = 1.2e-05, the median is 1.63 times higher in clients) between hsp90 clients and control nonclients when combining all disease phenotypes together.

### 7. Hsp90 clients have more pathogenic mutations compared to nonclients in the ClinVar dataset

ClinVar (https://www.ncbi.nlm.nih.gov/clinvar/) is a public archive aggregating the relationship between human variants and clinical status. According to the direct logic, we expect that hsp90 clients accumulate more often and more deleterious mutations because the hsp partially or completely compensates some or a majority of them. We used only nonsynonymous variants from ClinVar to check if our expectation is valid. We compared mutations in hsp90 client genes with all other genesandh the control list of nonclients (see above and Methods). We performed Fisher exact test to compare the number of pathogenic and benign mutations in hsp90 clients versus all other genes and versus control nonclients that are expected to have the same constraint level as well as the length. We observed an excess of pathogenic mutations in hsp90 clients versus all other genes (odds ratio = 5.17, p-value < 2.2e-16) as well as an excess of pathogenic mutations in hsp90 clients versus control nonclients (odds ratio = 5.96, p-value < 2.2e-16).

Next, we calculated the total number of mutations, the number and the proportion of pathogenic mutations, and the number of benign mutations per gene. We compared these numbers between hsp90 clients and nonclients (figure 4). We observed a significant difference in the total number of mutations, the number of pathogenic mutations, and the proportion of pathogenic mutations (Mann-Whitney U-test, p-value = 1.6e-06, 6.5e-09, 3.2e-06 respectively). The number of benign mutations per gene in hsp90 clients and nonclients was not significant (Mann-Whitney U-test, p-value = 0.18).

**Figure 4.**
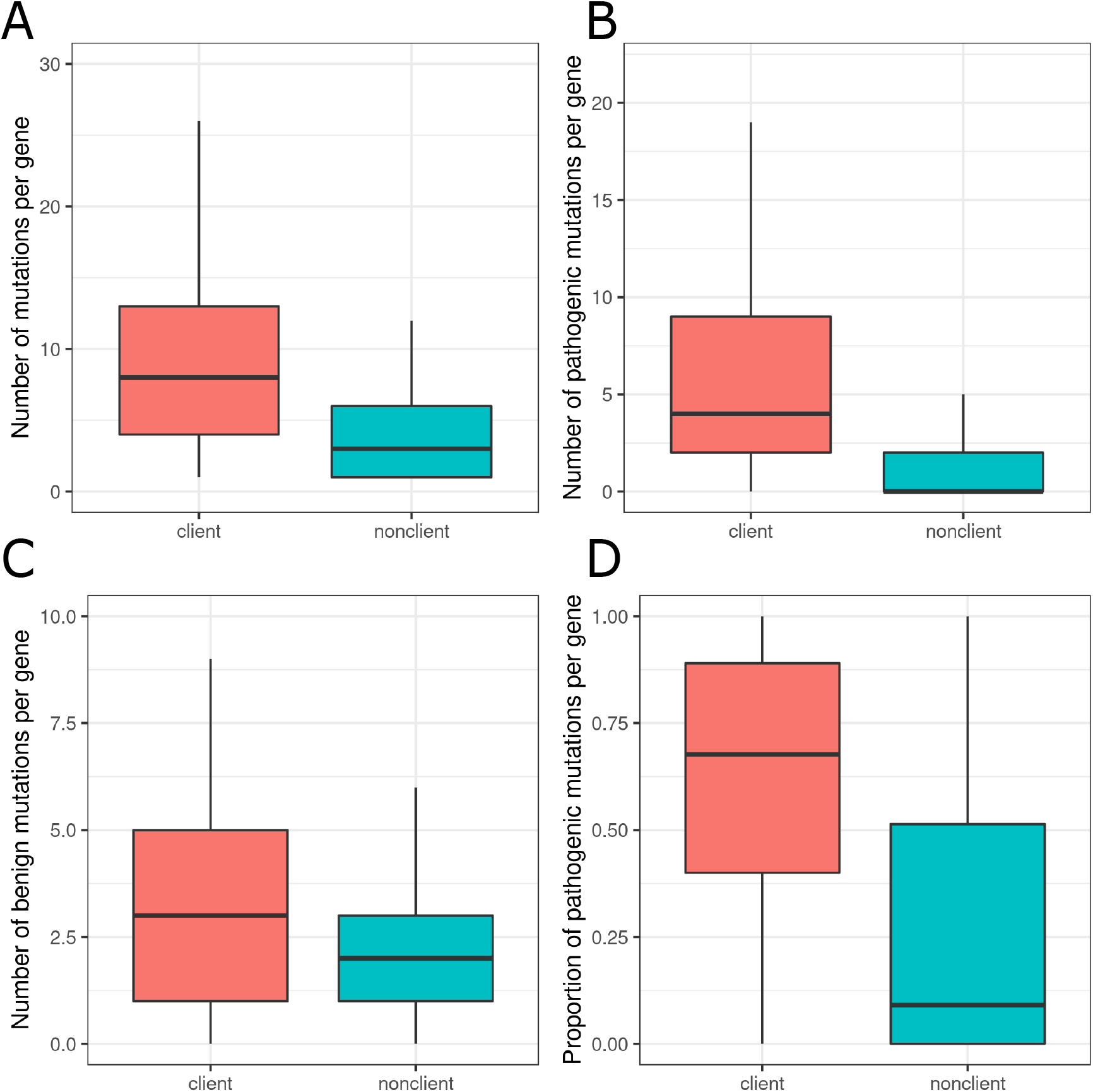
Comparison of the total number of mutations per gene (A), number of pathogenic mutations per gene (B), number of benign mutations per gene (C), and proportion of benign mutations per gene (D) in hsp90 clients and control nonclient genes. Data from ClinVar.

## DISCUSSION

In our work, using a spectrum of scenarios, assuming a buffering role of hsp90, we confirmed that hsp90 plays a more important role in species with low Ne, in tissues with increased cellular longevity and in genes directly affected by hsp90 (clients). All these findings are essential to take into account in the studies of ageing, molecular evolution and annotation of deleterious variants in the human genome.

## METHODS

1. Analysis of expression levels of hsp90 in primates We took expression data from (Brawand et al. 2011). In this work, RNA-seq data were generated for 10 animal species (Homo sapiens, Pan troglodytes, Pan paniscus, Gorilla gorilla, Pongo pygmaeus, Macaca mulatta, Mus musculus, Didelphis, Ornithorhynchus anatinus and Gallus domesticus) in 6 tissues (brain, cerebellum, heart, kidney, liver, testis). A set of constitutive aligned exons (perfectly aligned parts of genes) in each 1:1 orthologs group was built. We extracted data for 6 primate species on expression levels of hsp90 from supplementary data 1. Data about effective population size are from (Prado-Martinez et al. 2013) for grated apes species and from (Yuan et al. 2012) for Macaca.
2. Analysis of expression levels of hsp90 in human tissues To compare the expression level of hsp90 in human tissues with the different turnover rates we extracted expression data from Human Protein Atlas (https://www.proteinatlas.org/) and turnover rate data from (Mikhaylova et al. 2021) and (Sender and Milo 2021).
3. Comparison of the level of selection relaxation of hsp90 with a subset of close genes. To choose a subset of comparable with hsp90 genes (chapter 3) we performed Principle Component Analysis with gene-specific (for ∼11500 genes) characteristics of evolutionary constraints, such as
  - Residual Variation Intolerance Score - Residuals derived from the linear regression of the number of common functional variants on the total number of variants, the lower value, the more intolerant gene (Petrovski et al. 2013).
  - Probability of being loss-of-function intolerant - Posterior probabilities from a three-state Poisson mixture model, the higher value, the more intolerant gene (Cassa et al. 2017).
  - Selective effects for heterozygous protein-truncating variants (*SelCoefHet*) - Bayesian estimation of the selection coefficient on the heterozygous loss of function, the higher value the more intolerant gene (Cassa et al. 2017).
  - Functional Indispensability Score (FunInsScore) - Proteins are classified as ‘indispensable’ based on their impact on the network upon removal of the specific protein (Khurana et al. 2013).
  - Genome-Wide Haploinsufficiency Score (Haploins)- the more value, the more intolerant gene (Steinberg et al. 2015). As well as we added the following structural features of genes - Average exon length, GC content, Gene length, Number of transcripts, and Average number of exons per transcript (fig 3B). We used the first two principal components to calculate the euclidian distance of all genes to hsp90. We sorted all genes by this distance and chose the 300 closest ones for the following analysis. For each of the 1:1 ortholog groups of the chosen gene in mammals, we extracted species-specific values of the dN/dS ratio from Ensemb compara (Cunningham et al. 2021).
4. Choosing samples of clients and their control nonclients Client list were taken from(“Widespread Macromolecular Interaction Perturbations in Human Genetic Disorders” 2015). We performed Principal Component Analysis as in the previous paragraph but with additional metrics of evolutionary constraints (Karczewski et al. 2020):
  - oe-lof-upper-bin - Decile bin of LOEUF for given transcript (lower values indicate more constrained).
  - p - The estimated proportion of haplotypes with a pLoF variant. Defined as: 1 - sqrt(no_lofs / defined). We used this new PCA to analyse the distribution of clients of hsp90 in the space of the first two principal components and compare them with all other genes (chapter 5 in results). Also, we chose a control gene for each of the hsp90 client genes (chapter 4) as well as control genes for hsp90 itself (chapter 2).
5. Species-specific dN/dS calculation To calculate species-specific dN/dS values for hsp90, their control genes cops3 and ube3c respectively, as well as for their client and nonclient control genes we used a program codeml from PAML package v4.9 (Yang 2007)with following parameters - seqtype=1, model=2, NSsites=0, all other parameters were set as default. Codon alignments of each 1:1 ortholog group were taken from the OrthoMaM database v10c (Yang 2007; Scornavacca et al. 2019). A phylogenetic tree was taken from TimeTree (Kumar et al. 2022), which included >4000 mammals. Each ortholog group in codon alignment contained a different set and number of species which varied from 90 to 116 so accordingly the tree was pruned for each codon alignment to match the species sample.

## ACKNOWLEDGEMENT

The design of the experiment by KP and statistical analyses by VT were supported by Ministry of Science and Higher Education of the Russian Federation (agreement no. 075-15-2021-1084).

## REFERENCES

Agozzino, Luca, and Ken A. Dill. 2018. “Protein Evolution Speed Depends on Its Stability and Abundance and on Chaperone Concentrations.” Proceedings of the National Academy of Sciences of the United States of America 115 (37): 9092–97.

Alvarez-Ponce, David, José Aguilar-Rodríguez, and Mario A. Fares. 2019. “Molecular Chaperones Accelerate the Evolution of Their Protein Clients in Yeast.” Genome Biology and Evolution 11 (8): 2360–75.

Brawand, David, Magali Soumillon, Anamaria Necsulea, Philippe Julien, Gábor Csárdi, Patrick Harrigan, Manuela Weier, et al. 2011. “The Evolution of Gene Expression Levels in Mammalian Organs.” Nature. https://doi.org/10.1038/nature10532.

Cassa, Christopher A., Donate Weghorn, Daniel J. Balick, Daniel M. Jordan, David Nusinow, Kaitlin E. Samocha, Anne O’Donnell-Luria, et al. 2017. “Estimating the Selective Effects of Heterozygous Protein-Truncating Variants from Human Exome Data.” Nature Genetics 49 (5): 806–10.

Cunningham, Fiona, James E. Allen, Jamie Allen, Jorge Alvarez-Jarreta, M. Ridwan Amode, Irina M. Armean, Olanrewaju Austine-Orimoloye, et al. 2021. “Ensembl 2022.” Nucleic Acids Research 50 (D1): D988–95.

“HSP90 Shapes the Consequences of Human Genetic Variation.” 2017a. Cell 168 (5): 856–66.e12.

“HSP90 Shapes the Consequences of Human Genetic Variation.”. 2017b. Cell 168 (5): 856–66.e12.

Karczewski, Konrad J., Laurent C. Francioli, Grace Tiao, Beryl B. Cummings, Jessica Alföldi, Qingbo Wang, Ryan L. Collins, et al. 2020. “The Mutational Constraint Spectrum Quantified from Variation in 141,456 Humans.” Nature 581 (7809): 434–43.

Khurana, Ekta, Yao Fu, Jieming Chen, and Mark Gerstein. 2013. “Interpretation of Genomic Variants Using a Unified Biological Network Approach.” PLoS Computational Biology 9 (3): e1002886.

Kumar, Sudhir, Michael Suleski, Jack M. Craig, Adrienne E. Kasprowicz, Maxwell Sanderford, Michael Li, Glen Stecher, and S. Blair Hedges. 2022. “TimeTree 5: An Expanded Resource for Species Divergence Times.” Molecular Biology and Evolution 39 (8): msac174.

Marco, Moreno Di, Moreno Di Marco, Michela Pacifici, Luca Santini, Daniele Baisero, Lucilla Francucci, Gabriele Grottolo Marasini, Piero Visconti, and Carlo Rondinini. 2013. “Generation Length for Mammals.” Nature Conservation. https://doi.org/10.3897/natureconservation.5.5734.

Mikhaylova, A. G., A. A. Mikhailova, K. Ushakova, E. O. Tretiakov, V. Shamansky, A. Yurchenko, M. Zazhytska, et al. 2021. “Mammalian Mitochondrial Mutational Spectrum as a Hallmark of Cellular and Organismal Aging.” bioRxiv. https://doi.org/10.1101/589168.

Ohta, Tomoko. 1973. “Slightly Deleterious Mutant Substitutions in Evolution.” Nature 246 (5428): 96–98.

Petrovski, Slavé, Quanli Wang, Erin L. Heinzen, Andrew S. Allen, and David B. Goldstein. 2013. “Genic Intolerance to Functional Variation and the Interpretation of Personal Genomes.” PLoS Genetics 9 (8): e1003709.

Popadin, Konstantin, Leonard V. Polishchuk, Leila Mamirova, Dmitry Knorre, and Konstantin Gunbin. 2007. “Accumulation of Slightly Deleterious Mutations in Mitochondrial Protein-Coding Genes of Large versus Small Mammals.” Proceedings of the National Academy of Sciences of the United States of America 104 (33): 13390–95.

Prado-Martinez, Javier, Peter H. Sudmant, Jeffrey M. Kidd, Heng Li, Joanna L. Kelley, Belen Lorente-Galdos, Krishna R. Veeramah, et al. 2013. “Great Ape Genetic Diversity and Population History.” Nature 499 (7459): 471–75.

Rutherford, Suzanne L., and Susan Lindquist. 1998. “Hsp90 as a Capacitor for Morphological Evolution.” Nature 396 (6709): 336–42.

Scornavacca, Celine, Khalid Belkhir, Jimmy Lopez, Rémy Dernat, Frédéric Delsuc, Emmanuel J. P. Douzery, and Vincent Ranwez. 2019. “OrthoMaM v10: Scaling-Up Orthologous Coding Sequence and Exon Alignments with More than One Hundred Mammalian Genomes.” Molecular Biology and Evolution 36 (4): 861–62.

Sender, Ron, and Ron Milo. 2021. “The Distribution of Cellular Turnover in the Human Body.” Nature Medicine 27 (1): 45–48.

Steinberg, Julia, Frantisek Honti, Stephen Meader, and Caleb Webber. 2015. “Haploinsufficiency Predictions without Study Bias.” Nucleic Acids Research 43 (15): e101–e101.

Victor, Manish Prakash, Debarun Acharya, Sandip Chakraborty, and Tapash Chandra Ghosh. 2020. “Chaperone Client Proteins Evolve Slower than Non-Client Proteins.” Functional & Integrative Genomics 20 (5): 621–31.

“Widespread Macromolecular Interaction Perturbations in Human Genetic Disorders.” 2015. Cell 161 (3): 647–60.

Yang, Ziheng. 2007. “PAML 4: Phylogenetic Analysis by Maximum Likelihood.” Molecular Biology and Evolution 24 (8): 1586–91.

Yuan, Qiaoping, Zhifeng Zhou, Stephen G. Lindell, J. Dee Higley, Betsy Ferguson, Robert C. Thompson, Juan F. Lopez, et al. 2012. “The Rhesus Macaque Is Three Times as Diverse but More Closely Equivalent in Damaging Coding Variation as Compared to the Human.” BMC Genetics 13 (1): 1–12.

